# Rhythmic firing of neurons in the medulla of conscious freely behaving rats: rhythmic coupling with baroreceptor input

**DOI:** 10.1101/2022.03.04.483044

**Authors:** Bernat Kocsis, Irina Topchiy

## Abstract

Recent investigations emphasized the importance of neural control of cardiovascular adjustments in complex behaviors, including stress, exercise, arousal, sleep-wake states, and different tasks. Baroreceptor feedback is an essential component of this system acting on different time scales from maintaining stable levels of cardiovascular parameters on the long-term to rapid alterations according to behavior. The baroreceptor input is essentially rhythmic, reflecting periodic fluctuations in arterial blood pressure. Cardiac rhythm is a prominent feature of the autonomic control system, present on different levels, including neuron activity in central circuits. The mechanism of rhythmic entrainment of neuron firing by the baroreceptor input was studied in great detail under anesthesia but recordings of sympathetic-related neuron firing in freely moving animals remain extremely scarce. In this study we recorded multiple single neuron activity in the reticular formation of the medulla in freely moving rats during natural behavior. Neurons firing in synchrony with the cardiac rhythm were detected in each experiment (n=4). In agreement with prior observations in anesthetized cats, we found that neurons in this area exhibited high neuron-to-neuron variability and temporal flexibility in their coupling to cardiac rhythm in freely moving rats, as well. This included firing in bursts at multiples of cardiac cycles, but not directly coupled to the heartbeat, supporting the concept of baroreceptor input entraining intrinsic neural oscillations rather than imposing a rhythm of solely external origin on these networks. It may also point to a mechanism of maintaining the basic characteristics of sympathetic neuron activity, i.e. burst-discharge and cardiac-related rhythmicity, on the background of behavior-related adjustments in their firing rate.

## 1. Introduction

Cardiac rhythmicity is a hallmark of the autonomic control system from its sensory inputs all the way to sympathetic efferent nerve discharge directed to the heart and vasculature. However, whereas rhythmicity on sympathetic efferents is filtered before reaching peripheral targets ^1, 2^, on the afferent side it is reliably transmitted, with high fidelity. Rhythmic burst activity remains a functional component of sympathetic control of cardiovascular effectors ^1–4^, too, but that does not involve a direct oscillatory drive which is cut off by a low-pass filter system (~0.015 Hz for heart rate ^5^, and <0.5 Hz for vascular resistance ^6, 7^). Thus, it is the afferent side of this feedback loop, through the baroreceptor input, where neuronal networks may directly engage with oscillations intrinsically generated in the periphery, in the heart sinus node. The present study concerns the mechanism of how this rhythmic sensory input is integrated into the central rhythm in sympathetic networks. We tested in particular, whether the firing pattern of neurons in the medulla in conscious freely behaving rats is compatible with the concept ^8–10^ of the baroreceptor input entraining intrinsic neural oscillations rather than imposing a rhythm of solely external origin on these networks.

This concept was advanced on the basis of two lines of experimental evidence in anesthetized cats, i.e. (1) that the phase of rhythmic sympathetic bursts relative to the cardiac cycle changes with heart rate ^11^ and (2) that neurons in the medulla may fire rhythmically after barodenervation at frequencies overlapping with the cardiac rhythm but uncoupled from the heartbeat ^9, 12^. The original findings were later confirmed and extended to hundreds of neurons in different structures ^13–15^ and by extensive analysis of firing patterns using advanced mathematical techniques ^16–23^. With exception of a few studies in head-restrained cats ^24–26^ and guinea pigs ^27^ however, they all used anesthetized animals or decerebrate preparations, in acute experiments.

The anatomy of the medulla is not optimal for implantation of flexible microwires. It is relatively far from the dorsal surface, lying on the bottom of the fourth ventricle, and prone to minor dislocations or distortions associated with body movements or changes of intracranial pressure. Nevertheless, we demonstrate here that in freely moving rats, viable recordings can be made in these structures with stable spike activity, including neurons firing in synchrony with the cardiac rhythm. We used movable tetrodes mounted on microdrives which is by now considered a conventional technique to monitor multiple single neuron activity in behaving rodents. The medulla is the location of numerous structures participating in the control of sympathetic activity (see ^3, 28^ for the latest reviews). Just along the electrode track of our experiments (Fig. 1C), nuclei with widely different functions expressing cardiac rhythmicity ^13, 29–32^ are lined up. These include structures handling the baroreceptor input (solitary tract, NTS; Fig.1C) or regulating the sympathetic output in the spinal cord (ventrolateral medulla (RVLM), and neurons in the reticular formation integrating these nuclei into a complex network which extends to more rostral and caudal parts of the brainstem and is connected with higher order structures ^3, 28, 33–35^.

**Figure 1.**
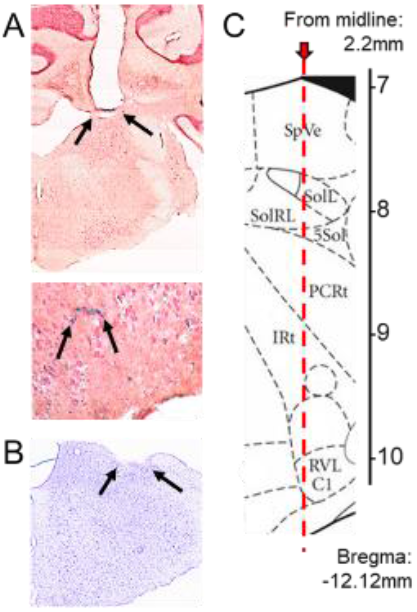
Localization of neurons recorded in the brainstem. **A**. Well-shaped tissue damage due to the electrode driving tube, dorsal to the medulla (*top*) and location of tetrodes in the ventral part of the medulla marked at the end of all recordings in that rat (*bottom*).**B**. Guiding tube at a deeper location in a different rat, potentially damaging sympathetic-related structures. **C**. Schematics of anatomical structures along the electrode track through the medulla.

In this study, we focused on neurons in the reticular formation, accessible in rats in the same coronal plane (intermediate and parvicellular reticular nuclei, PCRt and IRt; Fig. 1C) as NTS and RVLM and analogous with the primary structure exhibiting intrinsic 2-6 Hz oscillations in the cat ^13–15^. In agreement with observations in anesthetized cats, we found that neurons in this area exhibited high neuron-to-neuron variability and temporal flexibility in their coupling to cardiac rhythm in freely moving rats, as well. This included firing in bursts at multiples of cardiac cycles, but not directly coupled to the heartbeat. The data provided by our investigation is still far from sufficient to characterize this complex system however, which will require more chronic recordings in different types of neurons, in various brainstem structures, and in different behaviors. There may be species differences as well ^36–40^, relevant for translating the model to humans ^41^.

## 2. Methods

Male Sprague-Dawley rats were treated in accordance with National Institutes of Health guidelines. All experimental procedures were approved by Institutional Animal Care and Use Committee of Beth Israel Deaconess Medical Center.

Electrodes were implanted for chronic recordings of neuron spike activity in the medulla and peripheral cardiorespiratory signals to monitor the animal’s behavior and rhythmic activity in the cardiorespiratory system. Survival surgeries were conducted in sterile conditions under deep anesthesia, maintained by a mixture of Ketamine-Xylazine (i/p; 30-40 mg/kg ketamine and 5 mg/kg Xylazine) with supplementary injections of Ketamine (10 % of the initial dose) if necessary. Field potentials in forebrain structures were recorded with surface screw electrodes over the parietal cortex (n=3; AP: −3.5 mm, Lat: 2.5 mm from bregma) or with stainless steel wires implanted in the hippocampus (n=1; AP: −3.7, Lat: 2.2 mm, DV: −3.5 mm), muscle activity (electromyography, EMG) was monitored using multi-threaded electrodes in soft insulation for recordings in the neck muscle (n=2), and the diaphragm (n=1) and/or using the background activity on ECG electrodes (n=4). To recognize sympathetic-related spike activity, R-waves were detected in ECG or in diaphragmal EMG, with additional identification, where they were present, of cardiac cycles with longer R-R intervals, occurring regularly in many recordings, and occasional episodes of extrasystolia.

Recordings of the activity of neuron ensembles in the medulla were made using the procedures similar to those in our previous study ^42, 43^. For preparing the tetrodes, four polyimide insulated nickel-chrome microwires (12.5 micron diameter) were wound in a tetrode assembly station (Neuralynx, Inc) and bonded with heat gun. Three tetrodes were mounted in individually movable microdrives and assembled with a 18-pin connector (Omnetics; i.e. connecting to 12 tetrode wires and to 4 EMG, ECG, EEG electrodes, in addition to ground and reference) into a 2.6 gram covered headstage (Harlan 4 Drive, Neuralynx, Inc.). The tetrodes were lead through a guide tube placed above the medulla (AP −12.12 mm, Lat 2.2 mm, DV −8.0mm). All electrode wires and screws and the headstage with the microdrives were fixed to the skull with dental acrylic and all tetrodes were moved 1 mm out of the guide tube before the end of surgery. Electrophysiological recordings started after a 7-10 day recovery period.

Daily recording sessions lasted 2-6 hours during daylight period, in a 26×17×17cm recording box. After stable field potential, ECG, and EMG recordings were attained, the tetrodes were moved slowly (1/4 revolution, corresponding to 40 micron travel of the tetrode of the drive screw at a time, separated by 30-60 min) into the medulla until discriminable unit activity was found. The electrical signals were amplified, filtered (field potentials: 0.1-100 Hz, ECG and EMG: 0.1-3 kHz, units: 600-3 kHz), and sampled (16-bit, 10 kHz; Neuralynx, Inc.). The microelectrode location was marked at the end of the last recording by direct current, but its histological reconstruction was only successful in one experiment. The damage due to the tetrode guiding tube was clearly visible in postmortem histology in all experiments (Fig. 1) and thus the dorsoventral location of individual neurons along the electrode track was estimated relative to this marker by the number of turns of the microdrive.

Single neurons were identified and extracted off-line, based on their amplitudes and wave-shapes using principal component and K-means clustering algorithms (Spike2, Cambridge Electronic Devices, UK). Thus, the data points representing the spikes on four wires of a tetrode (Fig.3B) was reduced to 3 principal components which then were subjected to clustering. Principal Component Analysis (PCA) is a mathematical procedure that automatically extracts the features from data that contribute the most to the differences between the waveforms that make up the spikes. It is less susceptible to noise than other feature measurements and helps avoiding to make subjective decisions about the relevant features of the spike shape. The K Means algorithm is a well-known and fast clustering method of sorting spikes into clusters that are spherical and are of a similar size. The K-means algorithm follows an iteration sequence in which 3 steps of a cycle are repeated, i.e. a set of K cluster centers are defined, spikes are assigned to the nearest center, and the centers are recalculated. It should be noted that the K-Means algorithm always converges, but there is no guarantee that it converges to the optimum solution. Outliers were identified at more than 3 times the Mahalanobis distance from the center, and were eliminated. Units showing a refractory period of 2 ms or higher were considered as single units. Rhythmic components of spike trains and their relationship to (peripheral) cardiac rhythm were detected using time domain analysis methods. Spike counts were triggered by ECG R-waves or unit spikes to build auto- and cross-correlograms (Figs. 2,4,5) and raster plots were used to visualize the temporal dynamics of neuron activity (Figs. 5,6).

**Figure 2.**
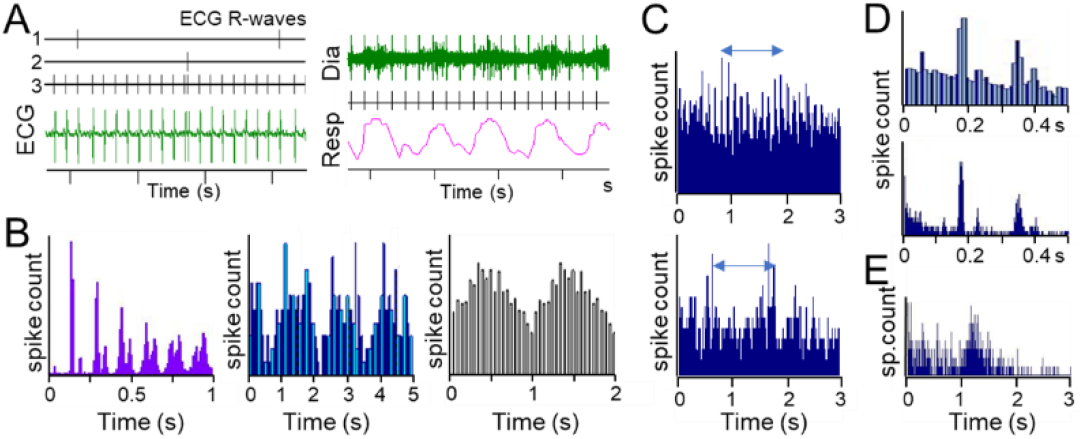
Identification of sympathetic related rhythmicity of medullary neurons. **A**. Peripheral signals to detect cardiorespiratory rhythmicity; *Left:* ECG R-waves to detect regularly occurring cycles with long R-R intervals (1) and rare episodes of extrasystoles (2) (cf. long R-R and double R in 3) and to monitor regular cardiac rhythm (3). *Right:* Diaphragmal EMG, to detect ECG R-waves, and respiratory rhythm. **B**. Examples of cross-correlograms of three neurons from three experiments, exhibiting cardiac rhythm (6 Hz, triggered by R-waves), slow oscillations (0.6 Hz, triggered by long R-R cycles), and respiratory rhythm (1 Hz). **C**. Slow oscillations detected on cross-correlograms triggered by regular R-waves, as modulation of the depth of cardiac peaks (*top*), corresponding to peaks in cross-correlograms triggered by long R-R cycles (*bottom*) (compare peak-to-peak times, marked by arrows). **D**. Cardiac rhythm in neuron autocorrelogram (*top*) and interspike interval histogram (ISIH, *middle*). Peaks correspond to analogous peaks in R-wave histograms (not shown). **E**.slow rhythm (Lf, 0.7 Hz) in ISIH (*bottom*). **B-E**y-axis: number of spikes per bin (normalized to peak to aid comparison of distribution patterns across histograms).

**Figure 3.**
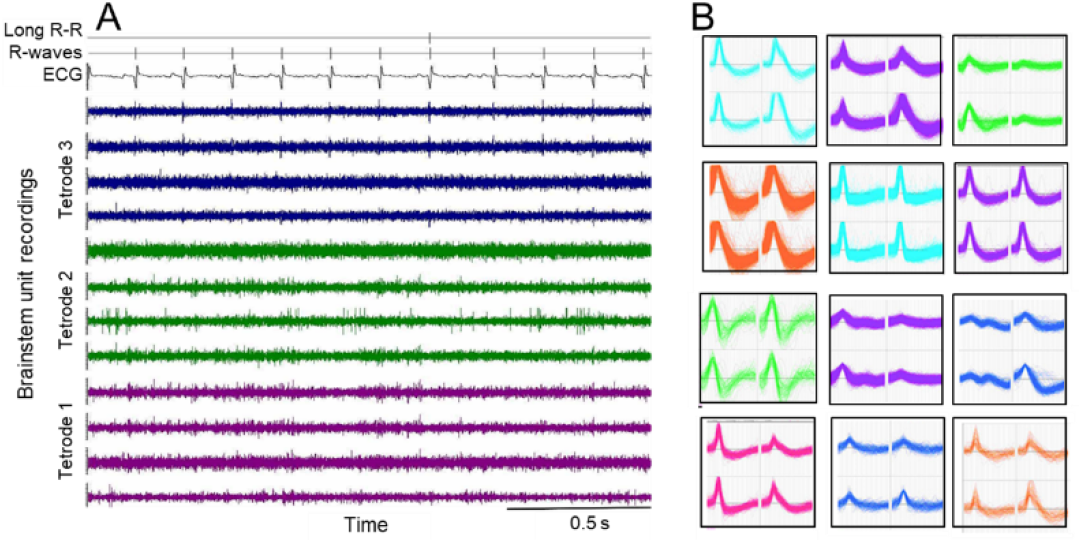
Identification of simultaneous activity of multiple neurons in three tetrodes in the medulla. **A**. Original sample recording of ECG with R-wave detection (regular and R-waves followed by long R-R interval) along with microelectrode recordings in the medulla (AP:-12.12, Lat:2.2, DV:8.75 mm, Fig. 1A) **B**. Spike shapes of 12 neurons as recorded on 4 electrodes of a tetrode, each, separated using standard spike sorting algorithms (see Methods).

**Figure 4.**
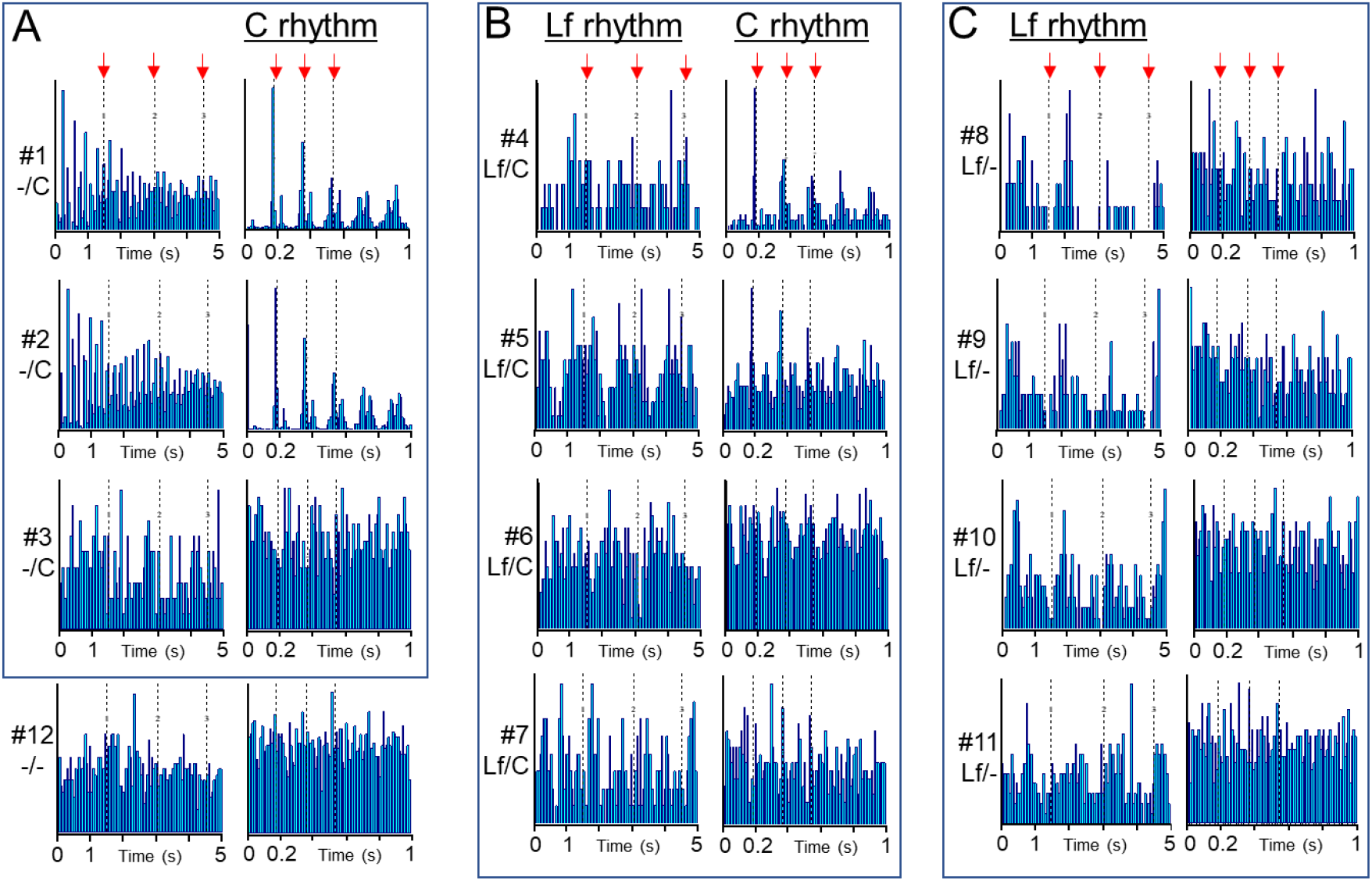
Variability of rhythmic coupling of single neuron activity in the medulla with cardiac activity (in the experiment also shown in Fig. 1A and 3). Crosscorrelogram of neuron firing triggered by regular R-waves or by cardiac cycles with long RR-intervals. Four neurons (#1,2,4,5) were identified on the first (Tetrode 1 in Fig.3), 5 neurons (#3,7,8,9,10) in the second, and 3 (#6,11,12) on the third tetrode. The neurons were classified depending on the predominant coupling expressed at heart rate (C rhythm, **A**), at low frequency (Lf rhythm, **C**) or both (**B**), as also labelled for each neuron, next to y-axis. Red arrows and dotted lines show the time of R-waves in the ECG autocorrelogram. y-axis: spike counts per time bin normalized to the largest peak; x-axis: time, note different time scales.

**Figure 5.**
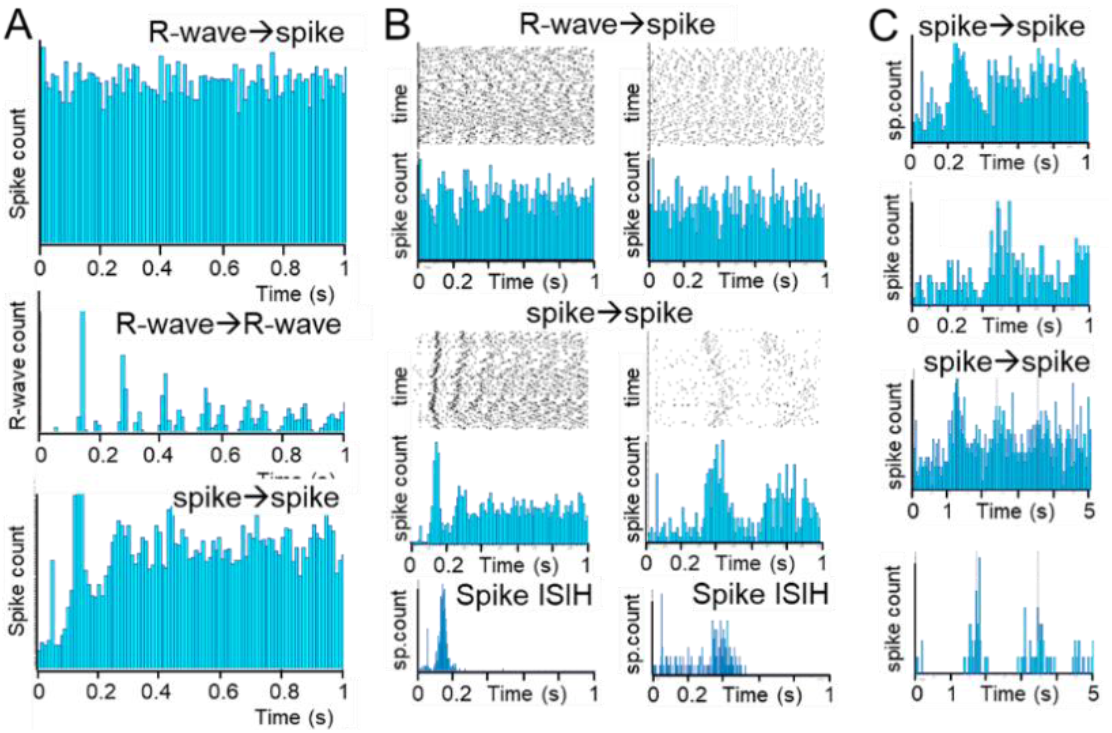
Cardiac rhythm in neuron activity with intermittent coupling with peripheral cardiac rhythm (for histology, see Fig.1 B). **A**. R-wave to spike crosscorrelogram (*top*)and autocorrelograms of R-waves (*middle*) and neuron firing (*bottom*) of a ~800 s long recording. **B**. R-wave to spike crosscorrelogram (*top*) and neuron firing autocorrelogram (*middle*), both with raster plot of spikes, and neuron interspike interval histogram (*bottom*) of a 65 s and a 60 s recording. **C**. Neuron spike autocorrelograms of several ~1 min long (70, 60, 80, 40s, from top to bottom) segments. Note prominent peaks in autocorrelograms in different segments at multiples of cardiac cycles (1 and 3 in B and at 2, 4, 8, and 12 cardiac cycles in C). Average spike counts per time bin were normalized to the largest peak in each diagram; note different time scales in C.

**Figure 6.**
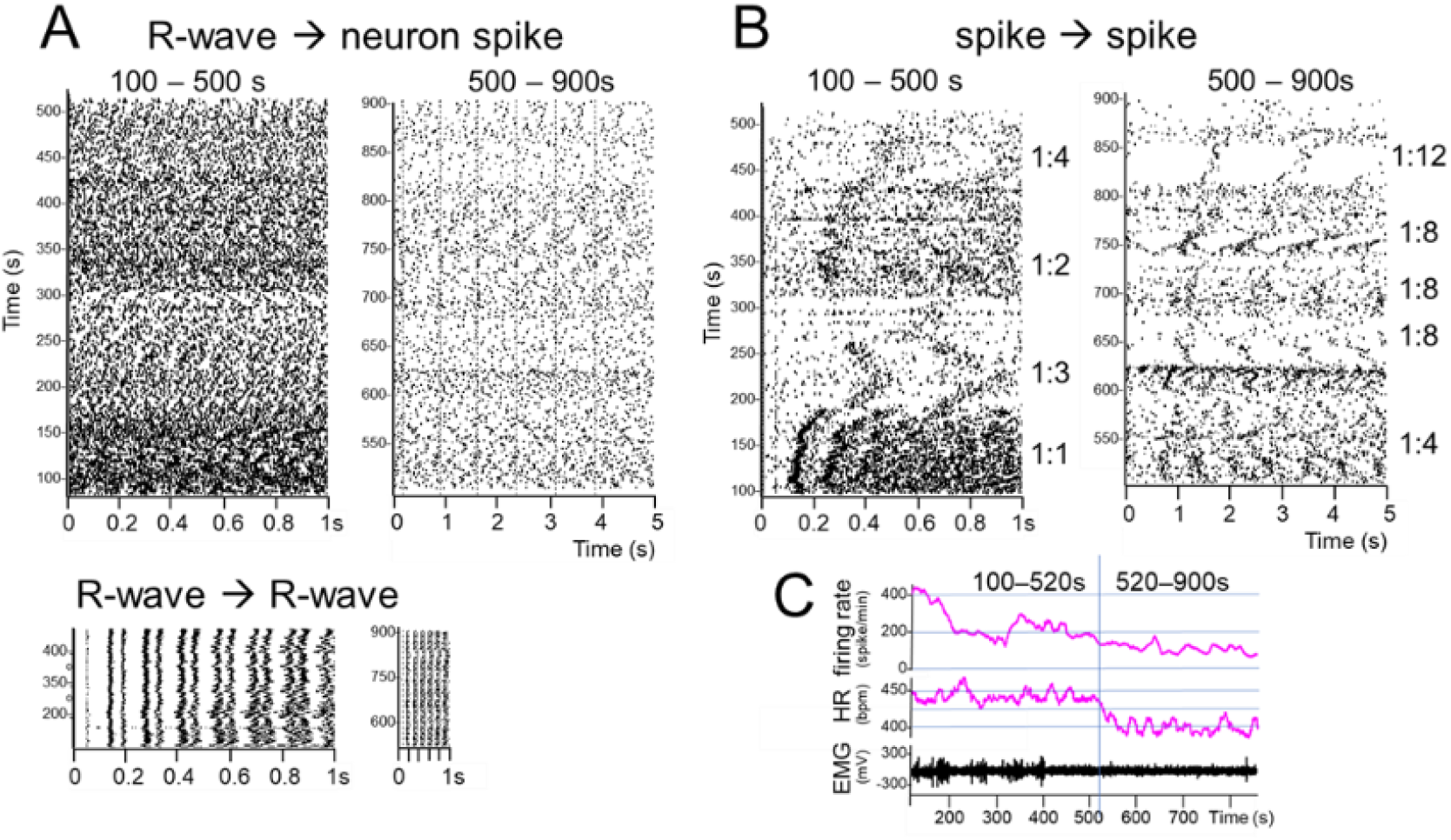
Time evolution of varying cardiac-related rhythm in the activity of a neuron in the reticular formation in the medulla (experiment also shown in Fig. 1B and 5) over a 800 s segment with a drastic decrease in heart rate at ~520 s and progressive decline in neuron firing rate. **A**. Neuron spikes (*top*) and ECG R-waves (*bottom*) triggered by R-waves in two consecutive segments (100 – 500 s and 500 – 900 s, note different time scales). **B**.Raster plot of the occurrence of neuron spikes triggered by neuron spikes of the same segments. Note drastic differences in firing rates between the two segments (*left* and *right* panels). **C**.Changes of firing rate (spikes per minute), heart rate (HR; beats per minute), and neck muscle EMG (mV).

## 3. Results

Recordings of multiple single neuron activity for this report were selected from neurons in the reticular formation (PCRt and IRt, Fig. 1C), i.e. below the NTS and above the RVLM, in 4 experiments. To recognize sympathetic-related spike activity, R-waves were detected in ECG or in diaphragmal EMG, with additional identification of cardiac cycles with longer R-R intervals, occurring regularly in every ~1.5-2 seconds in many (but not all) recordings, with occasional episodes of extrasystolic double R-waves (Fig. 2A). Respiratory rhythm was derived from diaphragmal EMG in one rat.

Characteristic sympathetic rhythms, synchronized with those in the periphery i.e. in ECG, were detected in medullary neuron activity, in all experiments. The frequencies varied in relatively narrow bands from one recording to the next. Thus, cross-correlograms, triggered by ECG R-waves or respiration, revealed three rhythms at different frequencies: cardiac rhythm (~5 Hz), a low frequency (Lf) oscillation triggered by long R-R cycles (0.5-0.7 Hz), and respiratory rhythm (1.2-2 Hz), as shown in the examples from three different rats in Fig. 2B. When long R-R cycles were not present, Lf rhythm could still be detected by other means, e.g. as rhythmic modulation of the depth of cardiac peaks in cross-correlograms triggered by regular R-waves (Fig. 2C) or Lf-related peaks in unit autocorrelograms and interspike interval histograms (Fig. 2E), along with C peaks (Fig. 2D).

Coupling of the activity of neurons in the medullary reticular formation to peripheral rhythms was highly flexible however. Expression of sympathetic rhythms in neuron firing varied in time and from one neuron to the next. Therefore, recordings of simultaneous activity of neuronal ensembles over time showed that neurons located in close vicinity could exhibit common rhythms in different combinations and at varying strengths and that coupling of the same neuron to sympathetic oscillations can drastically change in time. Next, we show two examples to demonstrate these two types of flexibility in oscillatory entrainment, using detailed analysis of spike trains in these recordings.

In the example shown in Fig. 3, simultaneous activity of 12 neurons was recorded on 3 tetrodes (Fig.3A). The neurons were identified by the shape of their action potentials as appeared on 4 electrodes of a tetrode (Fig.3B), using standard spike sorting algorithm (see Methods). Four neurons were identified on the first, 5 on the second, and 3 on the third tetrode. Spike counts in time bins, triggered by regular R-waves and by long R-R cardiac cycles (available in this experiment) were used to detect rhythmic components on different time scales (between 0-1s and 0-5s, Fig. 4). Cross-correlograms between ECG R-waves and neuron spikes revealed a within-group variability between closely packed neurons in coupling at heart rate (C) or at low frequency (Lf). Regular cardiac rhythm (i.e. “C”) was present in 7 neurons either alone (Fig. 4A) or together with Lf modulation (Fig. 4B). Lf rhythm could also be present alone with no C (Fig. 4C) or the spike trains could show no clear sign of either sympathetic rhythm (Fig. 4, cell #12).

The strength of rhythmic modulation of spike activity at different frequencies also varied from one neuron to the next. Compare for example strong C rhythm in neurons #1,2,4 with much weaker C in neurons #3,5,7 or strong Lf in cells #5,6,10 vs. weaker Lf in #7,11. When both present (Lf/C group, Fig.4B), either C or Lf oscillations could be dominant in different neurons; compare stronger C in cell #4, but stronger LF in #5 or similar C and Lf in cells #6,7. Furthermore, the delay of peak firing relative to R-wave differed between neurons, at both frequencies. In the cross-correlograms, C peak was coincident with R-waves in the majority of cells (#1,2,4,5) but could also appear at other phases of the cardiac cycle (#3,6,7). Time relationship between R-waves and neuron spikes at Lf also varied (compare for example cells #5 and 10).

In addition to “hard coupling” between cardiac rhythm and neuron firing revealed by R-wave triggered neuronal spikes shown in Fig. 4, pulse-related rhythmicity could manifest in the spike trains as intermittent coupling (sliding coordination) when the two rhythms (cardiac and neuronal) ran in parallel but were not phase-locked. Figures 5 and 6 show an example, in a rat with a potential “unintentional” barodenervation (unilateral, on the side of recording, see Fig.1B). In these neurons, R-wave triggered averaging will not show rhythmic activity, however strong peaks in the neuron autocorrelogram do appear at around the cardiac cycle (Fig. 5A). Unlike the stable rhythmicity in R-wave autocorrelogram however, the large peak in neuron autocorrelogram at the first cardiac cycle is followed by relatively minor elevations at multiples of the cardiac cycle barely standing out from the noisy “skyline” of background activity. However, in shorter (~ 1 min) segments, the neuron’s firing showed strong peaks in raster plots and in autocorrelograms (but not in R-wave to spike crosscorrelograms), at multiples of the cardiac cycle in a wide range from 1:1 to 1:12, alternating from one episode to the next (Fig. 5B, C). These episodes lasted about one min each, and occupied a total of 40% of the ~14 min-long recording.

Continuous monitoring of neuron firing using raster plots explains the origin of these peaks. Figure 6 shows the temporal evolution of this dynamic in a 13.7 min segment representing neuron activity in quiet waking (between time stamps from 100s to 500s, left panels in Figs.6A and B) and slow wave sleep (time stamps: 500s to 900 s, Figs.6A-B, right panels) with a sudden decrease in heart rate, relatively stable in both states (cardiac cycle 132 and 143 ms, respectively). The transition between the two states was also associated with a progressive decline in neuron firing rate (starting at ~400 s) and ceased motor activity (EMG; Fig. 6C).

In the first 3 min of the recording, the neuron fired rhythmically close to the heart rate (Fig. 6B, left). Starting at 170s however, the cycles of neuron firing were getting progressively longer and thus the phase between cardiac and neuronal rhythms started shifting. At 200s, the neuronal firing rate suddenly dropped and the cell fired at 3 times the cardiac cycle (see between 200 and 270s). The firing rate increased again at 320s and decreased at 450s. On this background, the probability of firing increased periodically at periods twice the cardiac cycle, and then (between 450-520s) slowed down to 4*132s (4*HR). This pattern of 1:4 coupling persisted in the beginning of slow wave sleep (Fig. 6B, right; note change in time scale on x-axis) but the period adjusted to the new, slower heart rate (see 520-600s). The firing rate kept fluctuating but at progressively lower rates, occasionally slipping into periodic firing at 8 times the cardiac cycle (640-800s), then for a short time (~1 min) at 12*143ms (at ~850s).

## 4. Discussion

The experiments, reported here, fill a major hiatus; as the rapid progress in technology in the past decades notwithstanding, we are only aware of few reports ^29, 30^ of neuron activity in central autonomic networks of the medulla exhibiting cardiac related rhythmicity, in conscious freely moving animals. In this study, we confirmed pulse-related activity in conscious rats indicating that rhythmic neuron firing in the medulla at characteristic frequencies of the cardiovascular system is not a property induced by anesthesia (or decerebration or head-restrain). Based on these results, we don’t see significant obstacles which would prevent large, comprehensive studies of central sympathetic networks in freely behaving animals, in the future.

Our study follows the extensive investigations of the complex system of autonomic networks in the medulla in prior studies that used anesthetized or decerebrate cats ^13–15^. These were recently extended to conscious animals (cats ^24–26^ and guinea pigs ^27^), but were still performed in acute experiments using head-restrain. We used the limited data set available from 4 conscious rats recorded in undisturbed conditions in this study to focus on a specific question of how rhythmic sensory input is integrated into the brainstem circuitry. For this, we chose recordings in the reticular formation, which is strongly involved in autonomic regulation, but owing to its diverse input and output connections is considered to play an integrative role in a variety of behaviors organized in the brainstem ^24, 44–48^ and to play a major role in bidirectional interactions with higher forebrain structures ^49–53^, including those marked with coupling between cortical oscillations and cardiac rhythm ^27, 54, 55^.

The pattern of neural activity in freely moving rats was found to adhere to the basic principles of organization proposed earlier ^8, 11, 23, 56^, based on recording from the reticular formation (lateral tegmental field) in anesthetized cats ^14, 15^, suggesting that cardiac-related rhythm in the reticular formation results from dynamic entrainment of neuronal oscillations by the sensory input ^10, 11^ rather than from a rhythmic drive unbendingly imposed on these neurons by the sensory input ^28, 33, 57^. The evidence supporting this model originally came from two sources: from experimental observations of rhythmic activity after barodenervation, uncoupled from the heartbeat, and from extensive analysis of this activity in the framework of coupled nonlinear oscillations. The first extended from early recordings (>50 years ago) of rhythmic sympathetic nerve discharge ^10, 58–60^to single neuron recordings in the brainstem ^12, 17^ in barodenervated animals (C and Lf also appear in human sympathetic fibers after bilateral baroreceptor deafferentation ^61^). The second line of evidence originated in early observations that the interval between the ECG R-wave and the cardiac-related bursts in sympathetic nerves ^11^ and central sympathetic neuron firing ^13^ was dependent on heart rate, as would be expected for a nonlinear oscillator, and was significantly reinforced later using advanced analysis techniques ^17–22^.

Our observations suggest that the central oscillatory circuits involved in sympathetic control exhibit strong flexibility of cardiac-related activity (shown in the examples of Figs. 4 and 5–6, in detail) in conscious rats, as well. Common oscillations, also present in the ECG, were detected in the firing of simultaneously recorded neurons in the medulla, but there was a variability within the network in the expression and dominance of C or Lf rhythms in different neurons and their phases relative to the ECG R-waves (Fig. 4). Systematic analysis of this latter in anesthetized cats revealed that spike to R-wave interval changed when the heart was paced at different frequencies ^13^ or when baroreceptor input was manipulated by changing the arterial blood pressure ^21, 62^. As arterial blood pressure and heart rate are essential components of cardiovascular regulation, this parameter will be of interest in future investigations using un-anesthetized animals to better understand autonomic control in complex behaviors, such as stress, exercise, arousal, sleep-wake states, etc.

Baroreflex and baroreceptor input itself is also a subject of behavior-dependent regulation ^63^. As shown under anesthesia, afferents from the carotid sinus, depending on their load, may promote different modes of coordination via flexible intermittent coupling with a central 2-6 Hz oscillator ^21, 62^. In this study, we also found neurons (Figs. 5–6) that may be equivalent to those reported in anesthetized or decerebrated cats, i.e. firing bursts in the characteristic frequency range of the cardiac rhythm but not necessarily phase-locked to the heartbeat ^9^. The origin of uncoupling of the neuron in Fig. 5–6 from the heartbeat remains unclear; it could be due to unintended barodenervation by damaging the NTS in this experiment (Fig. 1B). It could also be the consequence of blood pressure-dependent adjustments however, as shown under anesthesia ^21, 62^, indicating that oscillatory coupling in intact animals may dynamically change in different behaviors. Indeed, in a ~14 min recording of undisturbed behavior, in which the rat went through a state transition from quiet waking to slow wave sleep, the neuron shifted its activity to regular bursts at progressively increasing multiples of the basic 2-6 Hz rhythm. Importantly, this allowed the neuron to substantially decrease the rate of its firing while maintaining its basic characteristics, i.e. burst-discharge and cardiac-related rhythmicity. The implications of such intermittent rhythmicity in central sympathetic circuits on the dynamics of peripheral and central oscillations in baroreceptor intact animals will be an important topic of future studies.

The general concept of a rhythmic sensory input entraining central circuits may be considered for other feedback signals, as well. Respiratory rhythm, for example, generated in the brainstem ^64, 65^, reaches forebrain structures through sensory feedback from the periphery ^66–68^ where it selectively couples with the structure exhibiting local, intrinsic oscillations at matching frequencies in different behaviors, such as theta (6-10 Hz) in the hippocampus during sniffing and cortical delta (1-4 Hz) at rest ^67, 69, 70^, at least in rodents. Respiratory modulation of forebrain activity remains functional in humans as well, but due to a discrepancy between the frequencies of evolutionarily well-preserved cortical oscillations and a much slower respiratory rate (<0.5 Hz), it relies on other mechanisms, such as phase-amplitude coupling ^71^.

Heart rate in humans (>1 Hz) is within the frequency range of intrinsic oscillations of cortical networks, and the baroreceptor input reaches cortical structures, primarily the insula ^72–75^. There are reports of baroreceptor involvement in cognitive tasks and fear processing ^76, 77^ but the exact mechanisms remain poorly understood, lagging behind the recent progress in understanding the impact of the respiratory rhythm ^67^. Cardiovascular and respiratory control systems are closely intertwined on different levels of organization ^18, 78–83^ that besides their primary functions of supplying oxygen to tissues according to behavior might also impact their effects on higher brain functions.

## Acknowledgements

This work was supported by the National Institute of Mental Health grant R01 MH100820

## Author contributions

BK and IT contributed to the experiments, analysis, interpreting the results, and writing the report.

